# Probing the transcriptome response to shivering in skeletal muscle using a multilayered bioinformatics approach

**DOI:** 10.64898/2026.08.27.747657

**Authors:** Rosalie E. Baak, Guido J. E. J. Hooiveld, Patrick Schrauwen, Joris Hoeks, Reinier Raymakers, Anja van der Stolpe, Sander Kersten, Eric Kalkhoven

## Abstract

Cold acclimation holds therapeutic potential for improving metabolic health. We previously demonstrated that repeated cold-induced shivering enhances insulin sensitivity in humans. However, the molecular pathways that underlie the skeletal muscle shivering response, and how these relate to beneficial physiological effects, remain poorly understood. In this study, we combined complementary bioinformatics approaches to allow in-depth analysis of the transcriptomic response of human skeletal muscle to repeated shivering. We identified a robust transcriptional signature and show a sex-specific component in the shivering skeletal muscle response, which seemed to diminish following cold adaptation. Our findings provide mechanistic insights into cold-induced muscle adaptations, shed light on potential interesting molecular targets for further investigation, and emphasize the importance of including both sexes in future cold acclimation studies.

## Introduction

Skeletal muscle is part of the musculoskeletal system that provides form, support, stability, and movement to the human body. In addition, upon cold exposure, skeletal muscle plays an important role in maintaining core temperature by generating increased internal heat via two mechanisms: non-shivering thermogenesis (NST) and shivering thermogenesis (1,2). Whereas NST comprises heat production that is not associated with muscle activity, shivering thermogenesis involves rapid repeated skeletal muscle contractions, thereby leading to heat production through the inefficiency of ATP utilization (3). Interestingly, it has been shown that cold acclimation increases insulin sensitivity and that some level of muscle contraction is needed to provoke this effect (4,5).

More recently, we found that cold acclimation with shivering holds therapeutic potential in combating metabolic disease as it improved oral glucose tolerance, fasting glucose, triglycerides, non-esterified fatty acid concentrations and blood pressure (6). Notably, cold-induced shivering has been shown to stimulate the production and secretion of myokines that can exert beneficial metabolic effects on peripheral organs (7).

Previous analysis of the shivering transcriptome response by Sellers et al., 2024 revealed an upregulation of genes related to extracellular matrix (ECM) and cell adhesion genes indicative of muscle tissue remodeling, while gene sets linked to mitochondrial respiration were downregulated. Other studies have attempted to quantify the metabolic cost of shivering and fuel selection in humans (1,8–11). However, the molecular pathways that underlie the skeletal muscle shivering response and how these translate into beneficial physiological effects largely remain to be identified.

Obtaining significant and robust data from human cold physiology and in particular shivering studies is inherently challenging, since due to their typical resource extensive nature, study population sizes are generally limited. In addition, human interventions tend to be quite mild, creating further need for optimal approaches to dissect meaningful layers of biological information. To address these challenges, we combined various complementary bioinformatics approaches in an in-depth analysis of transcriptomic data from our previous cold acclimation study (6), which included both male (n = 11) and female (n = 4) participants. We used a combination of bioinformatics approaches to dissect meaningful layers of biological information within the shivering muscle transcriptome response (Figure 1), despite the biological variation observed. Together, these methods jointly assess the cold-induced gene network structure and capture individual-level variation, thereby extending beyond the limitations of the more commonly used group-level approaches.

**Figure 1.**
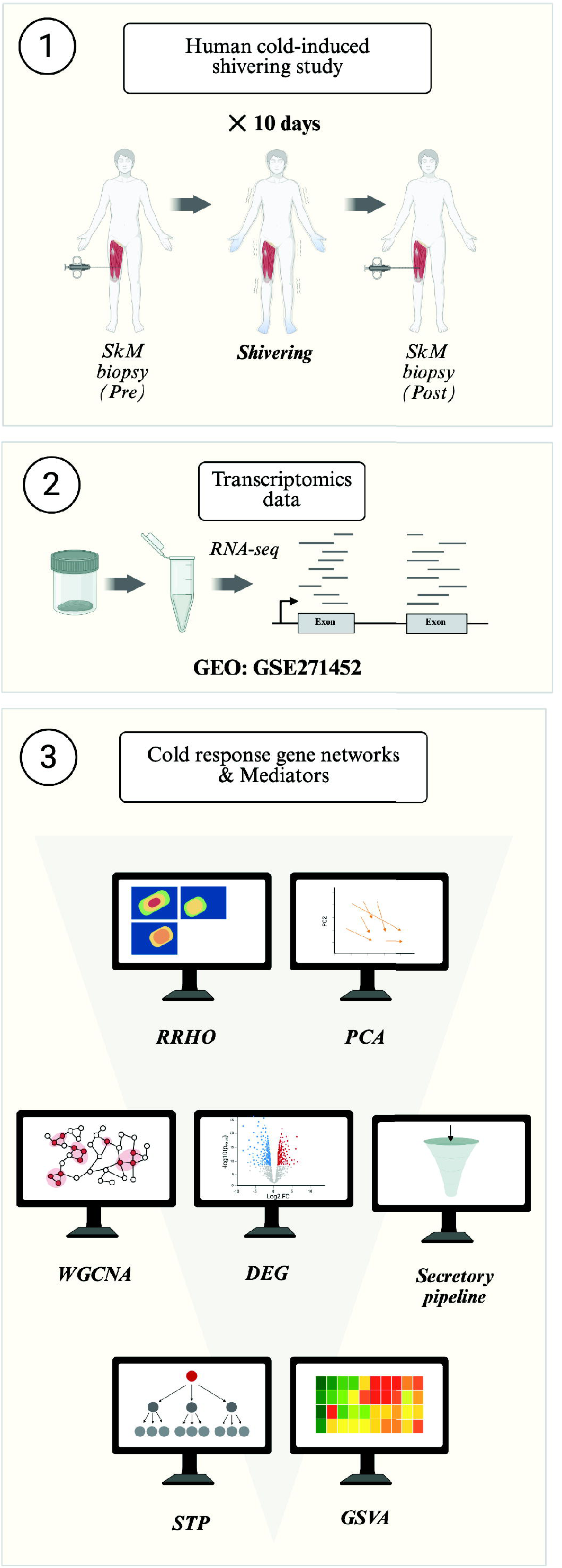
Schematic overview of the workflow to characterise the shivering muscle transcriptome response. We combined complementary bioinformatics approaches such as weighted gene co-expression network analysis (WGCNA), gene set variation analysis (GSVA), and signal transduction pathway (STP) analysis to dissect the shivering skeletal muscle transcriptome response, on the publicly available data from a previous performed cold acclimation study by Sellers *et al*., 2024

## Methods

### Source of data

The transcriptome data used for this paper were previously generated by our group and are available publicly: GSE271452 (6) and GSE156248 (4) (https://www.ncbi.nlm.nih.gov/gds). GSE271452 (platform GPL23227) comprises RNA-seq data from vastus lateralis skeletal muscle biopsies obtained from overweight/obese male and female individuals (body mass index 27–35 kg/m^2^) before and after cold acclimation (10 days, 1 hour of shivering per day). None of the subjects had diabetes. GSE156248 (platform GPL11532) comprises microarray data from vastus lateralis skeletal muscle biopsies obtained from male individuals with type II diabetes before and after 10 day cold acclimation (2h on day 1, 4h on day 2, and 6h on days 3 through 10). Both datasets employed a paired-sample design.

### Differentially expressed genes (DEG) identification

Differentially expressed genes were identified similarly for both datasets, described in detail by Sellers et *al.* (2024) by using generalized linear models that incorporate empirical Bayesian methods and defined as significantly changed when P ≤ 0.01 and fold change > 1.5. As a false discovery correction resulted in no differentially expressed genes, which is common for interventions in human subjects, we opted for a more lenient significance cut-off. Similar criteria were applied in a previous comparative transcriptome analysis involving human muscle biopsies PMID: 32887608. The lmFit and eBayes functions in the limma package (version 3.60.6) were used.

### Principal component analyses (PCA)

PCA was performed to reduce the dimensionality of a dataset while preserving as much of the variance (information) as possible. Transcriptome responses were visualised in 2D PCA space by arrows that connect pre-cold acclimation position to post-cold acclimation position for each participant. The pca and biplot functions from PCAtools (version 2.16.0) were used to do the analysis and visualise the first 2 principal components, respectively.

### Rank-Rank Hypergeometric Overlap (RRHO) analysis

RRHO analysis was performed to assess the similarity of the transcriptional responses observed in skeletal muscle after shivering cold acclimation (12) . The RRHO2 package (version 1.0) with default settings was used. We assessed the degree of differential expression overlap between data from Sellers et al. (2024) and data of a previous performed cold acclimation study by Hanssen et al. (2015) that also included shivering. Only genes that were common to both datasets were retained. Genes in each data set were ranked based on signed statistics (both log fold change and p-value). A 2D matrix of hypergeometric tests was then computed across all possible rank thresholds to identify regions with significant overlap. Resulting statistics of overlap were visualized in a heatmap, highlighting areas of concordant regulation between the two datasets.

### Network Analysis

The WGCNA package in R (version 1.73) was used to construct a co-expression network of genes (13) . In such a network, nodes are genes and edges represent the magnitude of co-expressing genes. WGCNA calculates the co-expression of genes based on an adjacency matrix, which in turn is derived from co-expression *similarity*. In WGCNA, the co-expression *similarity* (*s _ij_*) between two genes i and j, defined as the absolute value of the Pearson correlation coefficient:

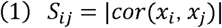

The Pearson correlation (inter-class) remains a valid method to measure co-expression of gene pairs in data from a paired design as it quantifies the relationship between two distinct variables (genes), rather than evaluating the reliability of repeated measurements of the same random variable, for which the intra-class correlation (ICC) is more appropriate (14).

Subsequently, the co-expression *similarity* (*s _ij_*) is transformed into the adjacency (*a _ij_*) using a soft-thresholding function:

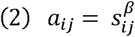

Where *β* ≥ 1 is a soft-thresholding power selected to approximate a scale-free topology as most biological networks exhibit scale-free properties. WGCNA then detects gene modules (densely interconnected *i.e.* co-expressed genes) by hierarchical clustering. The module eigengene (ME) of a module is defined as the first principal component thereby representing the overall expression level of the module. Highly connected genes within such a module are considered central regulators (i.e. hubgenes). Moreover, WGCNA assesses the association between modules and external metadata, thereby linking transcriptional signatures to clinical traits. To achieve the above, WGCNA calculates the gene significance (GS), measuring the correlation between gene expression and the external trait, and the module membership (MM), quantifying the connectivity of a gene within its module.

### Standard WGCNA with LMM

Li et al., 2018 previously reported an adapted strategy for WGCNA analysis of genomic data of paired design, which we applied here in our analysis. Firstly, we applied standard WGCNA to construct the co-expression network using a soft optimal threshold power of 4, corresponding to an R^2^ of > 0.8. The weighted network adjacency matrix was calculated which was then converted to dissimilarity measures. Subsequently, average linkage hierarchical clustering was used for gene module identification. Thirdly, modules that were at least 75% similar, were merged and assigned a unique color identifier. Importantly, to subsequently appropriately account for the correlation between paired samples from the same individual, we used the linear mixed-effects model to assess the association between the resulting gene modules and cold exposure and sex:

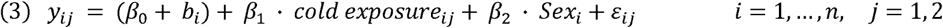

In the model, *y_ij_* is the expression level of the eigengene (i.e. the first principal component of a given module) of the i-th subject for the j-th condition. j = 1 is the pre-cold sample while j = 2 indicates the post-cold sample. Thus, cold exposure_1_ = 0 indicates baseline condition, and cold exposure_2_ = 1 indicates cold-exposed condition.

*β*_0_ is the overall intercept. *b_i_* the random effect that accounts for subject-specific baselines. *ε_ij_* is the residual error term. We assume that *b_i_* ~ *N*(*β*_0_, *τ*^2^), *ε_ij_* ~ *N* (0, *σ*^2^)and that *b_ij_* and *ε_ij_* are mutually independent. We used the test statistic and corresponding p-value derived from the linear mixed-effects model as measures of both strength and significance of the association between each gene module and phenotype/ cold exposure and sex (14) . Modules that were (1) specifically relevant to the pre-cold condition; (2) specifically relevant to the post-cold condition; (3) relevant to both cold condition and sex; and (4) specifically relevant to sex were evaluated. Then, gene significance for each node within these modules was defined as the absolute value of the test statistic from the mixed-effects model. Key hubgenes were identified for these gene modules by filtering on MM > 0.7, GS > 2.0 and restricting gene calls to initial assigned modules. Hubgenes were put into an overrepresentation analysis (ORA) for functional enrichment. Subsequently, hubgenes were tested for overlap with the DE genes and potential secretory proteins (described below). Overlapping genes were put into STRING network-based analysis for K-means clustering (15).

### Gene Set Variation Analysis (GSVA)

The GSVA package in R (version 1.52.3) was used to compute GSVA scores for GO Biological Process (GO- BP) gene sets based on the gene expression matrix (16) . Differentially active pathways were identified by using generalized linear models that incorporate empirical Bayesian methods and defined as significantly changed when P ≤ 0.01 and fold change > 1.5. The lmFit and eBayes functions in the limma package were used.

### Secretory pipeline

To identify potential secretory proteins, the 281 differentially expressed genes were first filtered for extracellular GO terms (GO:0005615 & GO:0005576), then screened for signal peptides using SignalP 6.0, and lastly filtered by average expression (logCPM ≥ 2), yielding 19 candidate secretory genes (S1 Table).

### Signal Transduction Pathway (STP) Analysis using STAP-STP (Simultaneous Transcriptome-based Activation Profiling of STPs)

STAP-STP technology (https://dcdc-tx.com) has been validated to enable quantitative measurement of pathway activity from transcriptomics data using Bayesian computational models (17,18). These models infer pathway activity scores based on the expression of 25–35 directly regulated target genes per STP (19–24). We applied STAP-STP to RNA-seq data of Sellers et al., 2024 (GSE271452) to analyze the activity of STPs. STP activity scores reported are quantitative measurements, normalized on a 0–100 scale. As each STP has a specific dynamic range, scores are not directly comparable between pathways. Differentially active pathways were assessed by using paired two-tailed t-tests and considered significantly changed when P < 0.05.

## Results

### The skeletal muscle shivering transcriptome response is robust despite inter-individual variability

We examined the transcriptome response in the vastus lateralis upon 10 days of cold acclimation with shivering to get a better molecular understanding of the impact of shivering on skeletal muscle (Fig1A and B). RNA-seq analysis was performed on muscle biopsies collected before and after cold acclimation in 15 subjects, which is very substantial for this type of invasive intervention. Pairwise comparison between these 2 conditions showed differential expression of 281 genes (227 upregulated, 54 downregulated; fold change > 1.5, p-value < 0.01).

First, we wished to assess the variability in cold acclimation and performed a principal component analysis (PCA) (Fig 2A). The PCA plot displays the pre-and-post conditions for the 15 participants along PC1 and PC2, with the transcriptome response in each participant indicated by grey arrows. Three key observations were made. Firstly, most participants show a response along the positive PC1-axis, which thus reflects cold acclimation. Secondly, the skeletal muscle transcriptomes clearly cluster on sex along PC2 (Spearman p -0.77, p-value < 0.01). And thirdly, the analysis revealed substantial variation in the transcriptome responses among participants, as reflected by the variation in both length and direction of the arrows. Thus, PCA reveals sex as a key source of variation within the skeletal muscle transcriptome and substantial inter-individual variation in the shivering transcriptome response. Patient characteristics at screening showed some variation but no overt sex-based skewing of metabolic parameters (S2 Table).

**Figure 2.**
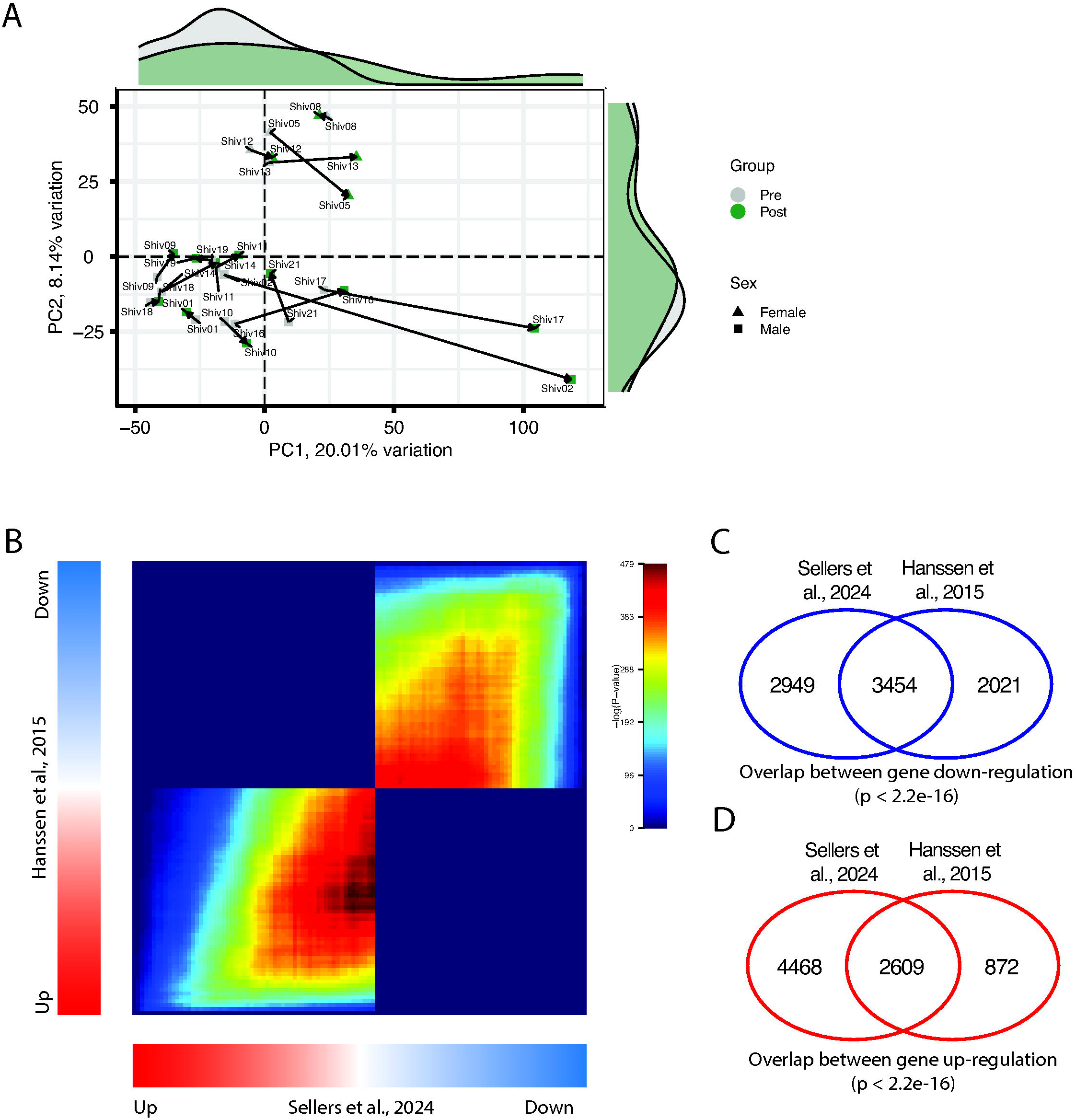
The skeletal muscle transcriptome response upon cold-induced shivering is robust despite inter-individual variability. A) Principal component analysis (PCA) of skeletal muscle transcriptomes before (grey) and after (green) cold acclimation of 15 participants (11 males = square, 4 females = triangle). Transcriptome responses are represented by arrows from pre-cold acclimation (base) to post-cold acclimation (head). B) Rank-rank hypergeometric overlap (RRHO) map comparing ranked gene lists by DE of two independent cohorts of cold studies, from vastus lateralus biopsies. Signal in the bottom left quadrant and upper right quadrant represent commonly upregulated and downregulated genes, respectively. C, D) Venndiagrams showing the overlap of genes that were both downregulated upon cold acclimation and that were both upregulated upon cold acclimation, respectively. Significance annotations for overlap indicate the results of Fisher’s Exact T-tests.

Together with this inter-individual variability identified by PCA, we also recognized similar transcriptional patterns between the studies of Sellers et al. (2024) and Hanssen et al. (2015) in response to cold exposure, particularly the cold-induced induction of ECM remodeling genes. Compared to Sellers et al. (2024), Hanssen *et al*., 2015 followed a milder cold-exposure protocol that did not lead to overt shivering. In addition, Hanssen et al. included subjects with type 2 diabetes, whereas Sellers et al. included subjects with overweight/obesity but not diabetes. Despite these differences and obvious limitations, we addressed potential similarities in transcriptional profiles between these two studies. To that end, RRHO was used (12) (Fig 2B). RRHO analysis revealed strong concordance along the diagonal, indicating similar up- and down-regulation of differential expression patterns of the studies. Interestingly, common downregulated genes (Fig 2C) were more prevalent than common upregulated genes (Fig 2D). We next investigated whether the top 20 most highly induced genes by the non-shivering cold protocol were also induced by the shivering protocol. Interestingly, the majority of the top 20 genes induced by the non-shivering cold protocol was also induced by the shivering protocol, despite the difference in study population, including many genes associated with the extracellular matrix (S3 Fig.). The gene expression changes were quite variable between volunteers, which is typical for human interventions.

Taken together, these findings show significant similarity in the gene expression profiles elicited by cold acclimation with and without overt shivering, despite differences in the study population.

### Network analysis reveals sex-biased network modules in shivering transcriptome response of skeletal muscle

Next, we aimed to further delineate potential shivering-regulated gene expression patterns. To investigate sex-biased transcriptional signatures and key mediators of the shivering skeletal muscle, we performed weighted gene co-expression network analysis (WGCNA) (13) (Fig 3A). Gene network construction resulted in the identification of 12 gene modules, each automatically assigned a unique color label by the package software. To subsequently relate these modules to metadata, a linear mixed effects model was used (for details see Materials and Methods). Fig 3B shows the association between the modules and two traits: cold condition (Post = 1, Pre = 0), and sex (Male = 1, Female = 0). Associations are shown as t-statistic resulting from the linear model and reflect both the strength and direction of the relationship. A positive t-statistic indicates higher module expression in the group encoded as 1 for the respective trait, while a negative t-statistic indicates high expression in the group encoded as 0.

**Figure 3.**
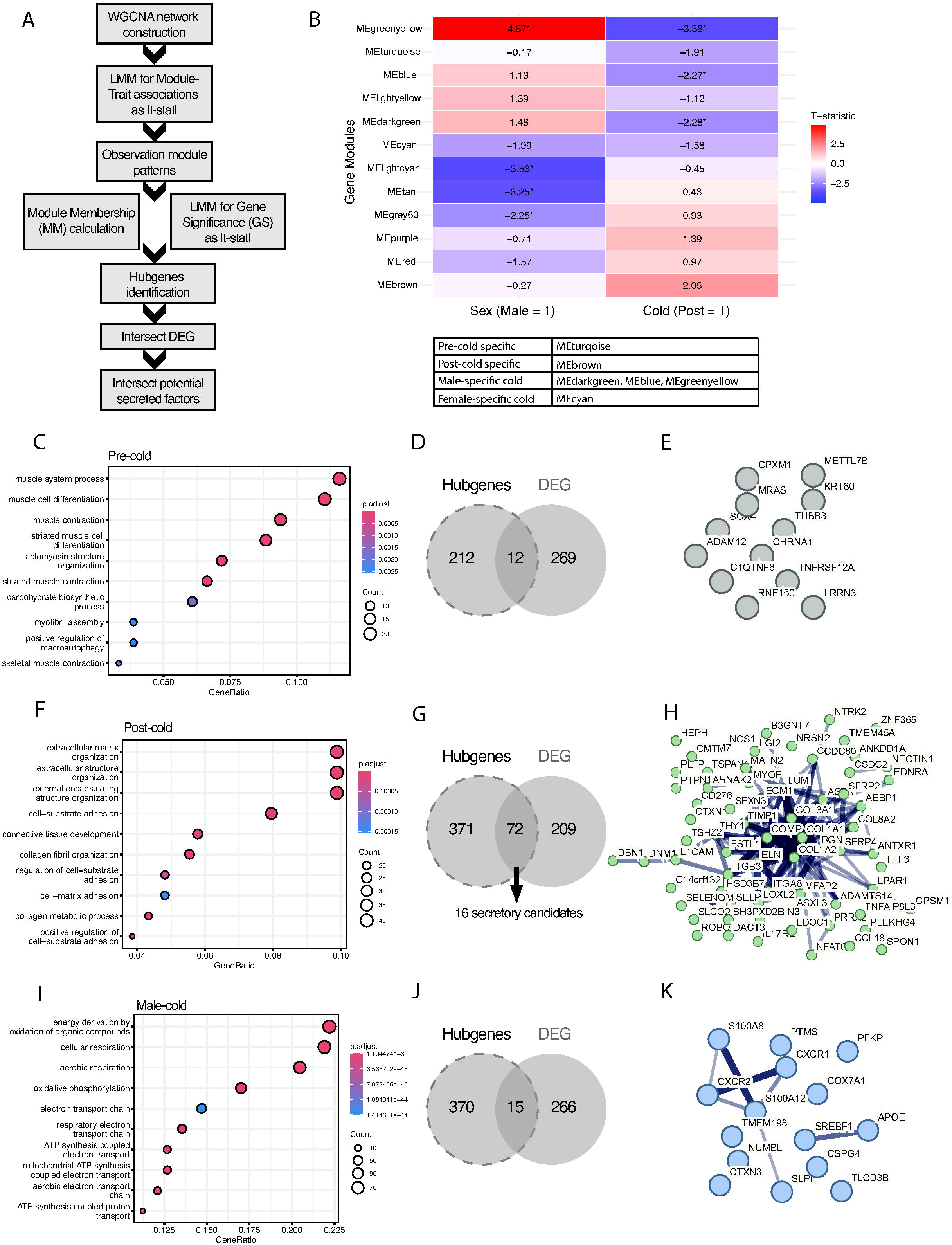
Network analysis reveals sex-biased network modules in transcriptomic cold response of skeletal muscle. A) Schematic representation of WGCNA pipeline. B) Heatmap of module-trait relationships for 12 identified modules, shown as absolute values of t-statistics resulting from the applied linear model. Appointed groups of modules are shown in the table on the right. C-E) Overrepresentation analysis of pre-cold specific, post-cold specific and sex-dependent cold response respectively. F-H) Venndiagrams of overlap of identified hubgenes with the DEG gene set. I-K) STRING PPi networks for the overlapping genes.

Overall, even though only a few women were included in the study, we observed a strong sex-dependent transcriptional pattern in skeletal muscle that is more pronounced than the effects of cold acclimation. This is reflected by both larger absolute t-statistic values and stronger statistical significances for module-sex associations compared to those for cold acclimation (Fig 3B). Despite being more subtle, the effect of cold acclimation reveals distinct module associations too, underscoring its physiological relevance. Given the mild nature of the intervention and inherent variability in human data, we chose to evaluate all modules with an absolute t-statistic of at least 2 for either sex or cold condition, regardless of statistical significance, to capture potential meaningful biological associations that may not reach conventional significance levels. Visual inspection of these module-metadata associations reveals (groups of) modules that were (1) specifically relevant to the pre-cold condition; (2) specifically relevant to the post-cold condition; (3) relevant to both cold condition and sex; and (4) specifically relevant to sex thereby reflecting basal sex-differences in SkM (Fig 3B below). The MEdarkgreen, MEgreenyellow and MEblue modules were associated with pre-cold acclimation (|t-stat| > 2, P < 0.05) and mildly with sex (|t-stat| > 1.13, P > 0.05) with higher expression in males (male-specific cold response). The MEcyan module associated with pre-cold acclimation (|t-stat| > 1.5, P > 0.05) with sex (|t-stat| = 2, P > 0.05) with higher expression in females (female-specific cold response). The MEturquoise module specifically associated with pre-cold condition (|t-stat| = 1.9, P > 0.05), and the MEbrown module specifically associated with post-cold condition (|t-stat| > 2, P > 0.05.). We also explored the module-cold association by comparing the Module significance (MS), which is defined as the mean GS across all genes in a module (S1A Fig.), and the relationships between GS and MM across all genes for each module that we investigated (S1B-G Fig.). The MEdarkgreen module had the highest relevance to the cold condition.

To identify potential key mediators in the shivering transcriptome response that may act in a sex-specific manner, we combined the modules described above with sex-biased expression and subsequently investigated encompassed hubgenes by overrepresentation analysis (ORA) for functional enrichment and testing overlap with DEG and potential secretory genes. We identified these secretory candidates (total 19) by filtering the DEG gene set for those associated with extracellular GO terms and possessing signal peptides (S1 Table). Firstly, the pre-cold specific module was enriched for hubgenes (224) that are related to basal muscle function (Fig 3C), which is consistent with expected baseline physiological activity. Twelve genes overlapped with the DE gene set (Fig 3D), but their protein products did not show functional connectivity in a protein-protein interaction (PPi) network from STRING (Fig 3E). This could suggest that some basal muscle genes are responsive to cold exposure but could act via distinct pathways. Secondly, the post-cold specific module was enriched for hubgenes (443) that are related to ECM remodeling (Fig 3F). Seventy-two genes overlapped with the DEG gene set (Fig 3G) and 16 of those were also secretory candidates. The protein products of these overlapping genes showed connections in a PPi network (Fig 3H). Notably, we identified Follistatin-related protein 1 (FSTL1) as a potential mediator in this module, an established glycoprotein secreted by skeletal muscle that has been associated with the promotion of endothelial cell function and angiogenesis in tissues (25,26). Additionally, potential mediators also included the cold shock domain-containing protein C2 (CSDC2), which is an RNA-binding factor that is induced by cold and is associated with regulation of mRNA stability (27,28). Thirdly, we identified 385 hubgenes across the three modules that exhibited male-biased expression and were more active at baseline, i.e. showing downregulation after cold acclimation. These hubgenes were enriched for cellular respiration and oxidative phosphorylation (Fig 3I). The higher baseline expression suggests a distinct metabolic state compared to females. Given the lower baseline expression of these modules in females, it is plausible that this pathway is less dynamically regulated in females, suggesting they may be less-relient on their modulation during cold acclimation. Consequently, the suppression of the oxidative phosphorylation-related genes may be a male-dominant regulatory response to cold acclimation. Fifteen genes overlapped with the DE gene set (Fig 3J) and their protein products showed some connectivity in a PPi network (Fig 3K). An interesting observation is the identification of Sterol regulatory element-binding protein 1 (SREBF1) as potential mediator in this module. SREBF1 is a key transcription factor that regulates the expression of genes that are involved in lipid homeostasis (29,30) and is described to regulate skeletal muscle cell size (29). Additionally, both CXCR1 and CXCR2 appeared in this module, which are both receptors for the IL-8 chemokine and play a crucial role in the recruitment of neutrophils to injured (31) and exercised (32) muscles.

We also identified 28 hubgenes from the module that showed female-biased expression in response to shivering that were slightly more active at baseline, i.e. showing downregulation after cold acclimation, as well as 95 hubgenes across 3 modules that exhibited female-biased expression at baseline, thereby potentially reflecting basal sex-differences in skeletal muscle. However, these genes showed no significant overrepresentation, nor overlap with the DEG gene set. This likely results from the underrepresentation of female samples (n = 4) in the present analysis.

Thus, WGCNA, like PCA, reveals that, in our shivering study, sex is a dominant factor shaping the transcriptional program of human skeletal muscle, which is already evident with only 4 women included in the study. Nonetheless, cold acclimation showed distinct gene module associations: while induction of ECM-related genes seems to represent a generic response, suppression of oxidative phosphorylation-related genes might reflect a male-biased adaptation. Although female-biased baseline and female-biased shivering-response mediators lacked significant overrepresentation likely due to underrepresentation of the number of female samples our data provide some evidence for a female-specific shivering response.

### Sex differences in skeletal muscle transcriptome are diminished upon shivering cold acclimation

To gain more insight into the molecular signaling pathways that are affected by shivering cold acclimation, and to assess sex differences at the pathway level, we applied both gene set variation analysis (GSVA) (16) and signal transduction pathway (STAP-STP) analysis (18). GSVA is a method that estimates pathway activity scores in individual samples based on expression data. Fig 4 shows a heatmap of the GSVA scores of the top 75 differentially active pathways upon shivering cold acclimation. Broadly, pathways upregulated upon shivering cold are predominantly involved in immune and inflammatory processes, lipid metabolism and musculoskeletal remodeling, whereas pathways downregulated upon shivering cold were predominantly involved in mitochondrial function, neuronal signaling and cellular stress responses. Interestingly, sample clustering shows a clear separation between males and females prior to cold exposure, while this separation seems attenuated post cold exposure. This implies that sex differences in the skeletal muscle transcriptome may be diminished upon shivering-induced cold acclimation.

**Figure 4.**
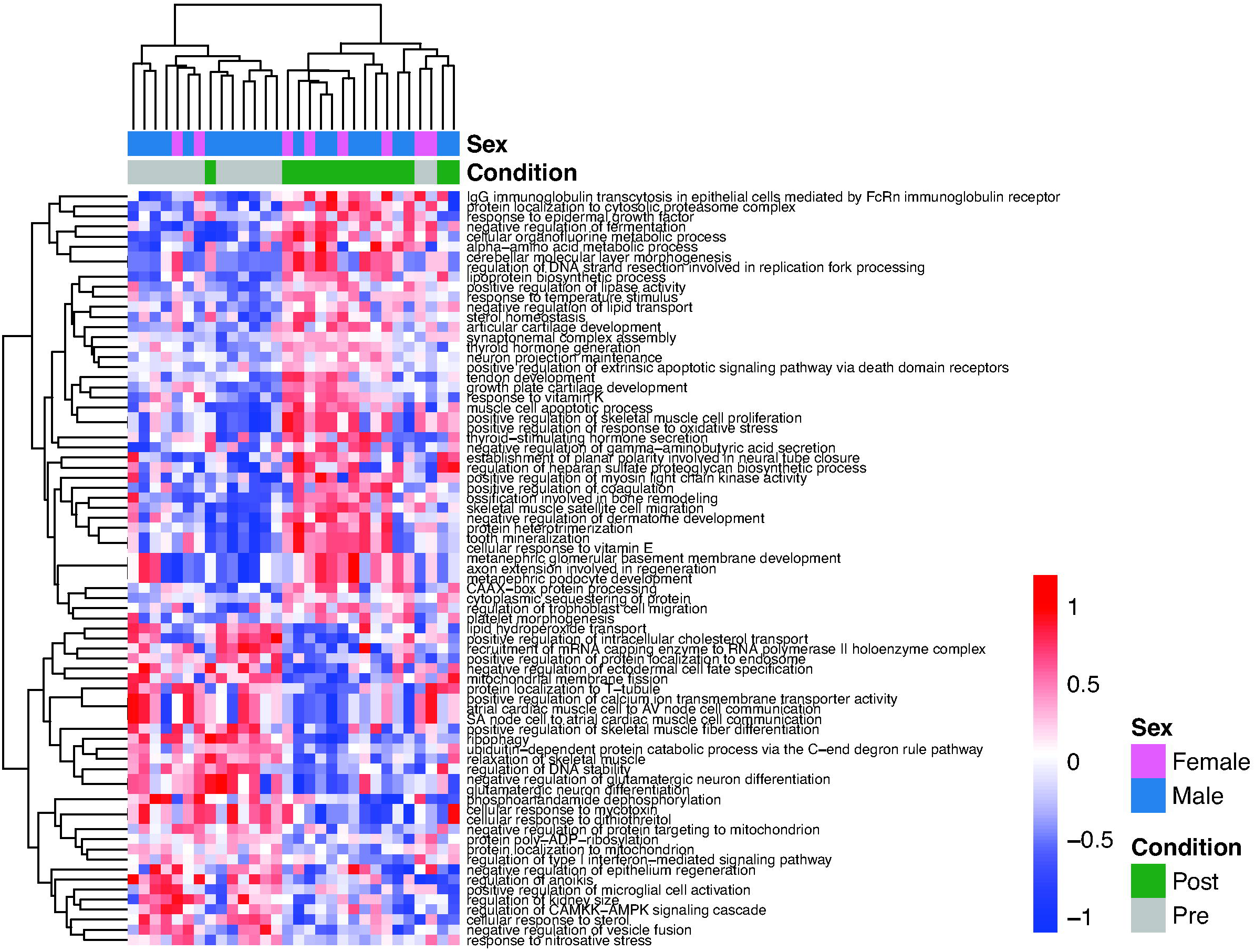
Sex differences in transcriptome of skeletal muscle are diminished upon shivering cold acclimation. Heatmap of top 75 differentially active pathways pre versus post cold acclimation, ranked on significance

STAP-STP is a knowledge-based computational method that infers the probability of signaling pathway activity in individual samples based on target gene expression data (18). Unlike conventional enrichment approaches, STP models actual pathway activity based on the expression of its downstream target genes rather than assuming a direct relationship between gene expression and protein activity. Application of STP to the shivering transcriptome data generated a heatmap showing pathway activities across the individual samples (S3 Table).

We analyzed 10 pathways and identified distinct sex-specific patterns of pathway activity for five of them. Particularly, shivering induced estrogen receptor (ER) activity specifically in males (Fig 5A), whereas shivering induced androgen receptor (AR) activity in both males and females (Fig 5C). Additionally, in males, shivering induced NFκB activity (Fig 5B), suggesting a more pronounced immune regulatory or inflammatory response in males compared to females. The PI3K-FOXO pathway serves to illustrate the specificity of this shivering response, as in this case females had baseline activity elevations, but its activity remained unchanged following shivering cold acclimation (Fig 5D). Interestingly, the shivering intervention appears to overall diminish most sex-based baseline differences in STP pathway activities. This emphasizes the convergence of shivering transcriptome responses between sexes as earlier observed in GSVA (Fig 4). In summary, our data provide evidence for a sex-specific skeletal muscle transcriptome responses upon shivering cold acclimation.

**Figure 5.**
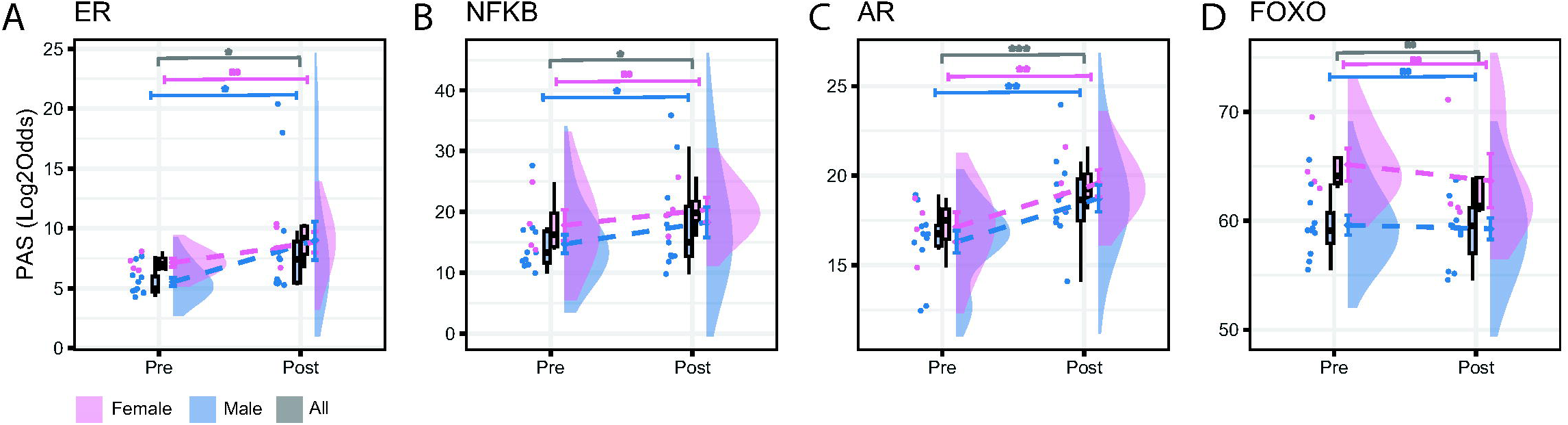
Shivering induces sex-specific skeletal muscle transcriptome responses. A-D) Raincloud plots depicting PAS (log2Odds) scores for ER, NFKB, AR and FOXO pathways, comparing pre- versus post-cold exposure. Each plot shows individual data points, half-violin plots representing score distributions, boxplots for group summaries, and connected group means ± standard error. Significance annotations indicate the results of paired t-tests comparing pre versus post scores across all participants (grey), as well as separately for males (blue) and females (pink). Asterisks denote levels of significance: * p < 0.05, ** p < 0.01, *** p < 0.001.

Additionally, both GSVA and STAP-STP methods that assess pathway activity for curated gene sets and molecular signaling pathways, respectively, suggest that sex differences in the muscle transcriptome are diminished upon shivering cold acclimation. Furthermore, STAP-STP analysis results provide molecular insight into the differentially affected signaling pathways upon cold-induced shivering.

## Discussion

Although cold-induced shivering has been attributed to contributing to beneficial physiological adaptations, the molecular pathways in skeletal muscle that underly these effects remain largely unexplored (33). The present study provides a framework for understanding the transcriptional adaptation of skeletal muscle upon shivering cold acclimation. Collectively, with a multilayered approach that combines complementary bioinformatics analyses, we uncovered a collection of genes that are regulated by cold-induced shivering in the human skeletal muscle, thereby identifying potentially interesting targets for further investigation. In addition, our results hint at a sex-specific component in the shivering skeletal muscle response, which seemed to diminish after cold adaptation.

Despite the relatively mild nature of the intervention and inherent challenges in human studies, our data reveal a robust and reproducible transcriptomic response to cold acclimation. Combined with the previously reported physiological benefits described by Sellers et al., 2024, these findings highlight the skeletal muscle sensitivity to cold-induced activation and underscore its potential as a target for prevention and treatment to metabolic disturbances. We find that shivering cold acclimation induces strong induction of gene expression while suppression of gene expression was more mild. We speculate that cold exposure triggers a cellular stress response that is characterised by rapid upregulated of genes driven by strong signalling cascades, whereas gene repression may occur through more subtle or indirect mechanisms, leading to comparatively modest fold changes.

We found that at baseline, male and female skeletal muscle exhibited distinct transcriptomic profiles. This finding aligns with previous studies that also demonstrated significant sex differences in the skeletal muscle transcriptome at baseline (34,35). STAP-STP analysis revealed distinct sex-specific responses to cold acclimation, which may reflect fundamental differences in adaptive strategies between males and females. Females had elevated baseline levels of hormonal signalling (ER pathway), immune regulation and ECM remodelling (TGFB pathway), and tissue maintenance (WNT pathway) (S3 Table). This distinct elevated transcriptional state in females might suggest that females might harbour more plasticity to condition an adaptive response. Males showed specific induction of estrogenic and NFKB STP responses, which could represent a compensatory mechanism potentially related to multiple processes involved in immune modulation, tissue remodelling or metabolic adaptations. Interestingly, although the skeletal muscle transcriptome is distinct between sexes at baseline and displays sex-specific responses to shivering cold acclimation, our data reveal a convergence of transcriptomic profiles following shivering cold acclimation, i.e. males and females are transcriptionally more similar post cold acclimation. This finding is similar to that of a previous endurance exercise study, where sex-based transcriptomic differences were also diminished after prolonged training (34).

The present study, which involved overt shivering, showed a marked effect of genes and pathways connected to extracellular matrix organization. These changes resemble the gene expression changes observed in a previous cold exposure study that was designed to avoid overt shivering (36). This similarity strengthens our conviction that the previous cold acclimation protocol, although designed to avoid overt shivering, might have actually triggered micro muscle contractions/shivering.

Overall, shivering cold acclimation and endurance exercise have been compared in the literature because they both involve repeated muscle contractions and require increased energy consumption (1). While multiple studies have investigated the effect of endurance exercise on the skeletal muscle transcriptome, few studies have examined cold acclimation. Nascimento et al., 2020 has demonstrated that the effects of exercise training and cold acclimation presumable involving microshivering on gene expression in human skeletal muscle are partially overlapping, yet remain distinct, which may be due to differences in motor unit recruitment, fibre type engagement, and contraction amplitude and frequency between cold-induced (micro) shivering and voluntary exercise (37).

Our study has several limitations. Firstly, the small sample size and particularly the low number of female participants limits our statistical power. Therefore, we urge caution for overrepresentation and emphasize the need for future studies with a balanced sex representation. It has been difficult to assess potential sex-based differences in the molecular adaptations to cold acclimation because most human cold physiology data are based on male subjects, with female physiology therefore being understudied. However, since significant sex differences at baseline (34,35) and upon endurance exercise training (34,38,39) have been reported, it is plausible that cold acclimation contains a sex-based component as well. Secondly, using unbiased complementary approaches and then intersecting them, we identified skeletal muscle candidate genes that may be regulated by cold-induced shivering, but these require functional validation. We acknowledge that transcript levels do not necessarily reflect protein abundance. Studies have assessed the relationship between transcript levels and protein abundance in exercised muscles and found that proteins with different functions are differentially regulated at the transcriptional level (40,41). Specifically, they found that exercise training induced the content of extracellular matrix-related proteins at the transcriptional level, while an increase in the content of mitochondrial proteins was not thus might be controlled by post-transcriptional mechanisms. This phenomenon has yet to be investigated for repeated muscle contractions induced by shivering cold acclimation.

Therefore, proteomic profiling of skeletal muscle upon shivering cold acclimation could provide valuable information on which processes are in effect altered on the protein level. Lastly, skeletal muscle is well established as a secretory organ that produces and releases cytokines and peptides into the circulation, which are collectively termed myokines (42–46). These myokines can exert autocrine, paracrine or endocrine effects, thereby influencing a range of systemic physiological processes. To explore this aspect, we implemented a secretory pipeline to identify potential cold-induced secretory proteins. Notably, Popp et al., 2025 introduced MultiSTEP, a method to assess the impact of genetic variations on secreted proteins, and found that almost half of missense variants impact secretion, post-translational modifications (PTM), or both (47). This highlights another layer of complexity in the interpretation of cold-induced systemic effects.

Understanding shivering response mechanisms holds therapeutic potential in combating metabolic disease (33). Our analysis of the skeletal muscle transcriptome response allowed us to shed light on skeletal muscle candidate genes that may be important for transducing the metabolic beneficial effects of cold acclimation reported earlier (6). We expect future studies to benefit from and expand upon on our work because our study touches upon considerations for future protocol design for human cold interventions as well as shedding light on potentially interesting targets for further investigation.

## Declarations

### Ethics approval and consent to participate

Not applicable

### Consent for publication

We hereby provide consent for the publication of the manuscript detailed above.

### Availability of data and materials

The analysed data in this manusript are the publicly available datasets GSE271452 (6) and GSE156248 (4), which were sourced from Gene Expression Omnibus (GEO) database.

### Authors contributions

JH, PS, SK, AS, RR, and EK designed and supervised the study; REB, GJEJH, AS, and SK performed data-analysis; REB performed dry-lab; REB, GJEJH, AS, SK, and EK interpreted results; REB prepared figures; REB and EK drafted the manuscript; All authors reviewed the manuscript and approved the final version of the manuscript.

## Acknowledgements

The authors thank Dzhansel Hashim and Denis P. Blondin for helpful discussions and support.

## Supporting information

**S1 Fig. Gene significance for cold exposure status across gene modules.**

A) Barplot showing the mean gene significance across all genes per gene module resulting from linear mixed model testing cold exposure. B-G) Relationships between gene significance and absolute module membership across all genes per gene module.

**S2 Fig. Top 20 upregulated genes during Hanssen et al., 2015 cold exposure compared to Sellers et al., 2024 shivering cold exposure.**

Heatmap of top 20 upregulated genes pre versus post cold acclimation in Hanssen et al., 2015, for all individuals of both Sellers et al., 2024 and Hanssen et al., 2015.

