## Supplementary material for "Probing the transcriptome response to shivering in skeletal muscle using a multilayered bioinformatics approach"

**Supplementary information:**

Supplementary table 1: secretory candidate genes

Supplementary table 2: Participant screening characteristics

Supplementary table 3: STP activity profiles

Supplementary figures 1 and 2

**Supplementary table 1. Nineteen shivering-induced sercretory candidate genes**

| EnsembleID | GeneName | AveExpre | log2FC | pvalue |
|---|---|---|---|---|
| ENSG00000108821 | COL1A1 | 5,80170556 | 1,7339661 | 0,00026246 |
| ENSG00000100979 | PLTP | 3,35513276 | 0,89501988 | 0,00036803 |
| ENSG00000049540 | ELN | 3,16839733 | 1,30456748 | 0,00161478 |
| ENSG00000182492 | BGN | 4,55686142 | 1,3423419 | 0,0020789 |
| ENSG00000168542 | COL3A1 | 6,16017777 | 1,16534813 | 0,00212856 |
| ENSG00000179796 | LRRC3B | 2,22344202 | -0,6566039 | 0,00224373 |
| ENSG00000163430 | FSTL1 | 5,36918517 | 0,94138142 | 0,00349363 |
| ENSG00000106819 | ASPN | 2,2805519 | 1,06832652 | 0,00349934 |
| ENSG00000173546 | CSPG4 | 4,18735247 | 0,63329748 | 0,00364571 |
| ENSG00000091986 | CCDC80 | 4,2143979 | 1,00506675 | 0,0037436 |
| ENSG00000164692 | COL1A2 | 6,32081117 | 0,97828047 | 0,00393397 |
| ENSG00000154096 | THY1 | 2,60847773 | 1,50238074 | 0,00503249 |
| ENSG00000130203 | APOE | 3,49716252 | 0,60037977 | 0,00506016 |
| ENSG00000139329 | LUM | 3,84805917 | 1,07975923 | 0,00604408 |
| ENSG00000106624 | AEBP1 | 3,32614906 | 1,46984319 | 0,00628203 |
| ENSG00000143369 | ECM1 | 2,95589655 | 0,58761097 | 0,00854488 |
| ENSG00000106483 | SFRP4 | 2,80011638 | 0,99774928 | 0,00882595 |
| ENSG00000102265 | TIMP1 | 3,60768214 | 0,93799601 | 0,00906542 |
| ENSG00000134013 | LOXL2 | 2,02604979 | 1,14027046 | 0,00967958 |

The 281 DEG were filtered for extracellular GO terms GO:0005615 and GO:0005576, then screened for signal peptides using SignalP 6.0, and lastly filtered by average expression (logCPM ≥ 2), yielding 19 candidate secretory genes

**Supplementary table 2 Participant characteristics at screening**

| Participant characteristics | Mean± SD | | |
|---|---|---|---|
| | Total[1] (n=15) | Males (*n*=11) | Females (*n*=4) |
| Age (years) | 62 ± 7 | 62 ± 7 | 63 ± 6 |
| Height (cm) | 174.4 ± 9.2 | 179 ± 0.1 | 163 ± 0 |
| Body mass (kg) | 92.7 ± 14.5 | 97.7 ± 13.6 | 79.0 ± 4.3 |
| BMI (kg/m$^2$) | 30.3 ± 2.9 | 30.6 ± 3.1 | 29.7 ± 2.6 |
| Systolic blood pressure (mmHg) | 138 ± 11 | 142.3 ± 7.4 | 129.6 ± 9.3 |
| Diastolic blood pressure (mmHg) | 88 ± 8 | 89.4 ± 7.3 | 83.7 ± 4.2 |
| Fasting plasma glucose (mmol/L) | 5.5 ± 0.5 | 5.5 ± 0.5 | 5.4 ± 0.3 |
| Fasting serum insulin (pmol/L) | 79.2 ± 68.2 | 92.6 ± 75.8 | 42.5 ± 10.8 |
| Haemoglobin (mmol/L) | 9.1 ± 0.8 | 9.4 ± 0.7 | 8.4 ± 0.5 |
| HbA1c (%) | 5.4 ± 0.3 | 5.4 ± 0.3 | 5.3 ± 0.2 |

Data are presented as mean± s.d. BMI, body mass index; HbA1c, glycated haemoglobin.
[1)] As reported previously in Sellers et al., 2024.

**Supplementary table 3. STP activity profiles pre versus post cold acclimation**

| | Subject | Sex | FOXO | MAPK | AR | ER | HH | NFKB | NOTCH | STAT1/2 | TGFB | WNT |
|---|---|---|---|---|---|---|---|---|---|---|---|---|
| Pre | Shiv10 | Male | 55,5 | 6,5 | 18,9 | 5,0 | 7,2 | 13,4 | 16,1 | 12,9 | 23,5 | 5,0 |
| | Shiv11 | Male | 65,6 | 4,7 | 18,7 | 5,9 | 6,9 | 17,4 | 11,9 | 12,0 | 20,8 | 6,1 |
| | Shiv12 | Female | 63,0 | 5,1 | 17,9 | 6,7 | 7,1 | 18,1 | 16,8 | 17,1 | 21,8 | 8,5 |
| | Shiv13 | Female | 63,6 | 3,5 | 18,7 | 7,3 | 7,0 | 14,5 | 19,7 | 17,1 | 22,5 | 10,6 |
| | Shiv14 | Male | 59,1 | 2,6 | 16,7 | 4,6 | 7,0 | 11,1 | 9,1 | 8,1 | 16,2 | 8,4 |
| | Shiv16 | Male | 56,3 | 2,4 | 15,9 | 4,3 | 7,1 | 17,0 | 15,8 | 13,3 | 20,0 | 3,5 |
| | Shiv17 | Male | 61,7 | 4,2 | 12,4 | 7,7 | 7,1 | 27,6 | 13,8 | 31,1 | 23,4 | 10,6 |
| | Shiv18 | Male | 59,3 | 1,2 | 16,3 | 5,5 | 7,2 | 11,9 | 14,6 | 8,3 | 17,1 | 10,2 |
| | Shiv19 | Male | 57,0 | 3,0 | 17,2 | 4,8 | 7,1 | 9,9 | 11,1 | 15,4 | 18,9 | 6,2 |
| | Shiv01 | Male | 59,1 | 4,4 | 17,2 | 4,7 | 7,1 | 12,2 | 8,3 | 11,1 | 22,8 | 3,4 |
| | Shiv21 | Male | 59,8 | 8,9 | 16,5 | 7,5 | 7,1 | 16,7 | 12,3 | 18,2 | 25,2 | 9,1 |
| | Shiv02 | Male | 63,4 | 4,7 | 16,7 | 6,1 | 6,6 | 13,3 | 35,6 | 13,4 | 19,6 | 10,2 |
| | Shiv05 | Female | 69,5 | 4,8 | 17,0 | 6,5 | 6,9 | 13,8 | 13,9 | 15,9 | 23,7 | 11,3 |
| | Shiv08 | Female | 64,5 | 5,0 | 14,9 | 8,1 | 7,1 | 24,9 | 13,3 | 27,3 | 24,2 | 9,6 |
| | Shiv09 | Male | 58,9 | 2,5 | 12,7 | 4,8 | 7,1 | 11,3 | 10,4 | 9,3 | 21,0 | 5,4 |
| Post | Shiv10 | Male | 59,6 | 5,3 | 19,5 | 7,5 | 7,1 | 15,0 | 17,7 | 13,5 | 22,3 | 5,3 |
| | Shiv11 | Male | 63,7 | 6,1 | 20,8 | 7,4 | 6,9 | 19,5 | 13,5 | 14,1 | 20,5 | 11,1 |
| | Shiv12 | Female | 60,8 | 4,6 | 19,6 | 6,7 | 7,1 | 19,4 | 15,1 | 12,7 | 24,8 | 9,3 |
| | Shiv13 | Female | 61,5 | 13,1 | 21,6 | 10,1 | 7,0 | 15,9 | 15,7 | 17,5 | 26,5 | 9,3 |
| | Shiv14 | Male | 55,2 | 2,4 | 17,2 | 5,5 | 7,0 | 12,9 | 17,1 | 11,7 | 19,2 | 2,4 |
| | Shiv16 | Male | 58,7 | 5,2 | 17,4 | 9,8 | 7,2 | 22,3 | 23,1 | 16,8 | 25,9 | 6,3 |
| | Shiv17 | Male | 59,0 | 22,8 | 18,9 | 20,4 | 8,6 | 35,9 | 40,6 | 33,8 | 25,8 | 5,9 |
| | Shiv18 | Male | 62,3 | 1,2 | 18,6 | 5,4 | 7,1 | 12,6 | 10,2 | 8,1 | 18,7 | 9,0 |
| | Shiv19 | Male | 55,4 | 0,9 | 17,9 | 5,4 | 7,1 | 9,8 | 7,9 | 11,2 | 20,1 | 4,9 |
| | Shiv01 | Male | 54,6 | 2,3 | 17,6 | 5,3 | 7,1 | 11,8 | 18,5 | 11,8 | 23,0 | 7,7 |
| | Shiv21 | Male | 63,9 | 7,7 | 24,0 | 8,0 | 7,1 | 17,9 | 26,7 | 14,8 | 21,9 | 11,2 |
| | Shiv02 | Male | 59,5 | 37,9 | 20,1 | 18,0 | 8,9 | 30,7 | 33,1 | 31,3 | 28,2 | 4,7 |
| | Shiv05 | Female | 71,1 | 18,0 | 19,0 | 10,4 | 5,7 | 25,7 | 17,0 | 23,3 | 31,9 | 8,4 |
| | Shiv08 | Female | 61,2 | 5,5 | 18,1 | 8,2 | 6,9 | 20,4 | 17,3 | 25,5 | 22,8 | 11,2 |
| | Shiv09 | Male | 60,1 | 3,1 | 14,1 | 5,8 | 7,0 | 12,8 | 7,3 | 8,6 | 19,5 | 10,9 |

**Two-sided paired t-test**

**Pre/post - All**

| | FOXO | MAPK | AR | ER | HH | NFKB | NOTCH | STAT1/2 | TGFB | WNT |
|---|---|---|---|---|---|---|---|---|---|---|
| Statistic | -0,80 | 1,89 | 4,56 | 2,80 | 0,74 | 2,34 | 1,76 | 1,13 | 2,22 | -0,02 |
| P-value | 0,43 | 0,08 | 0,00 | 0,01 | 0,47 | 0,03 | 0,10 | 0,28 | 0,04 | 0,98 |
| Significance | ns | ns | *** | * | ns | * | ns | ns | * | ns |

**Pre/post - Males**

| | FOXO | MAPK | AR | ER | HH | NFKB | NOTCH | STAT1/2 | TGFB | WNT |
|---|---|---|---|---|---|---|---|---|---|---|
| Statistic | -0,32 | 1,35 | 3,35 | 2,44 | 1,40 | 2,27 | 1,80 | 1,17 | 1,45 | 0,10 |
| P-value | 0,75 | 0,21 | 0,01 | 0,03 | 0,19 | 0,05 | 0,10 | 0,27 | 0,18 | 0,92 |
| Significance | ns | ns | ** | * | ns | * | ns | ns | ns | ns |

**Pre/post - Females**

| | FOXO | MAPK | AR | ER | HH | NFKB | NOTCH | STAT1/2 | TGFB | WNT |
|---|---|---|---|---|---|---|---|---|---|---|
| Statistic | -1,39 | 1,70 | 6,55 | 1,78 | -1,19 | 0,75 | 0,17 | 0,15 | 1,76 | -0,44 |
| P-value | 0,26 | 0,19 | 0,01 | 0,17 | 0,32 | 0,51 | 0,87 | 0,89 | 0,18 | 0,69 |
| Significance | ns | ns | ** | ns | ns | ns | ns | ns | ns | ns |

**Female/Male - Pre**

| | | | | | | | | | | |
|---|---|---|---|---|---|---|---|---|---|---|
| Statistic | 3,18 | 0,66 | 0,80 | 3,21 | 0,12 | 1,05 | 0,55 | 1,65 | 2,27 | 2,84 |
| P-value | 0,02 | 0,52 | 0,45 | 0,01 | 0,91 | 0,34 | 0,59 | 0,15 | 0,04 | 0,02 |
| Significance | * | ns | ns | ** | ns | ns | ns | ns | * | * |

**Female/male - Post**

| | | | | | | | | | | |
|---|---|---|---|---|---|---|---|---|---|---|
| Statistic | 1,65 | 0,36 | 0,80 | -0,06 | -1,73 | 0,64 | -1,04 | 0,97 | 1,94 | 2,17 |
| P-value | 0,18 | 0,73 | 0,45 | 0,96 | 0,14 | 0,53 | 0,32 | 0,36 | 0,12 | 0,05 |
| Significance | ns | ns | ns | ns | ns | ns | ns | ns | ns | * |

Table contains Pathway Activity Scores (PAS) across individual samples on a normalized scale (see Methods). Color coding ranges from blue for most inactive to red for most active. Statistics are shown below and indicate the result of two-sided paired t-tests comparing pre versus post scores across all participants, as well as separately for males and females, and male versus female both at baseline and post cold acclimation. Asterisks denote levels of significance: *p <0.05, ** p < 0.01, *** p <0.0001. Note: For Pre vs Post tests, t-statistics are computed as Pre – Post. For Male vs Female tests, t-statistics are computed as Female – Male. Thus, a positive t-statistic indicates higher values in the first group.

**Supplementary Figure 1. Gene significance for cold exposure status across gene modules.**

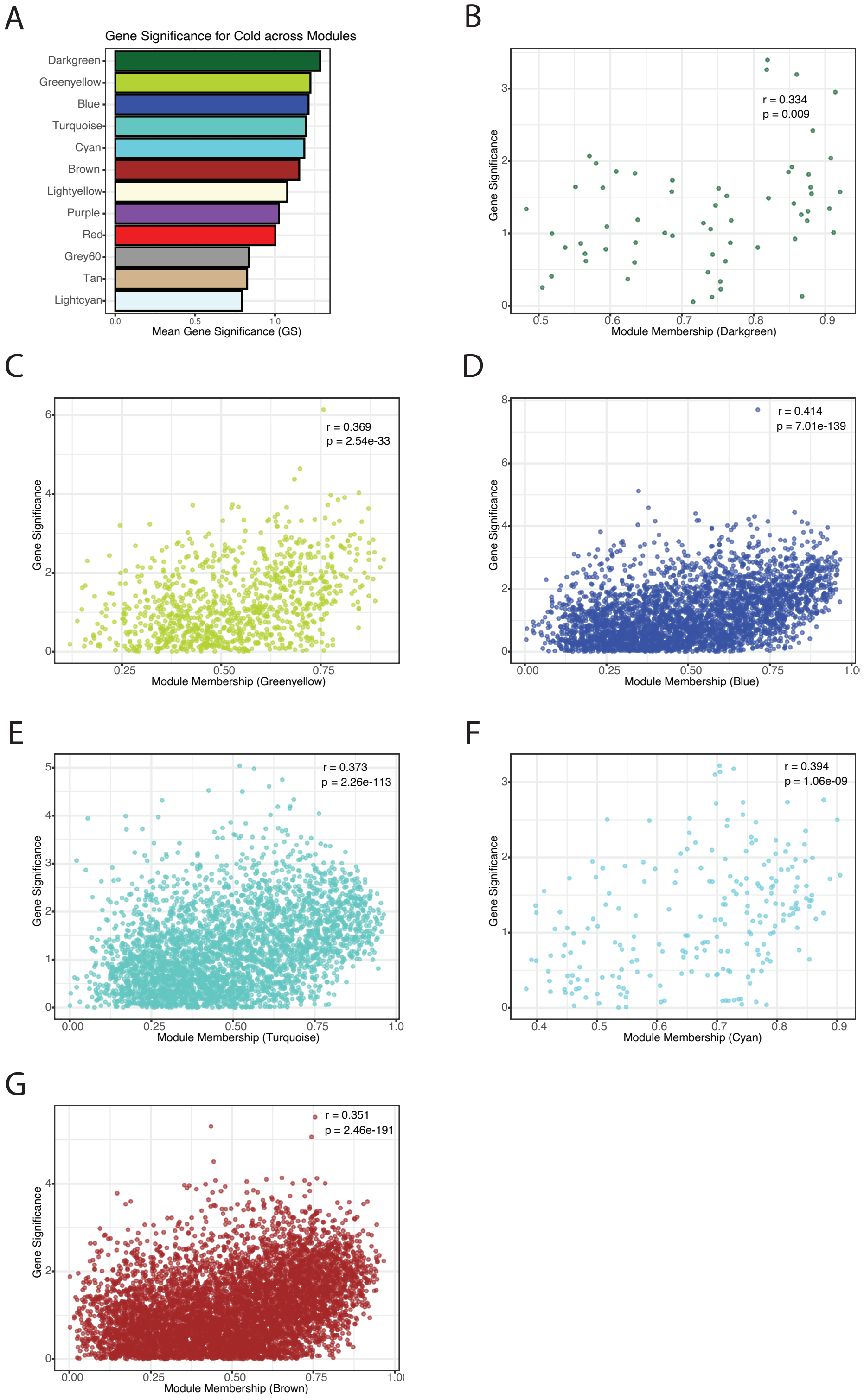


A) Barplot showing the mean gene significance across all genes per gene module resulting from linear mixed model testing cold exposure. B-G) Relationships between gene significance and absolute module membership across all genes per gene module.

**Supplementary Figure 2. Top 20 upregulated genes during Hanssen et al., 2015 cold exposure compared to Sellers et al., 2024 shivering cold exposure.**

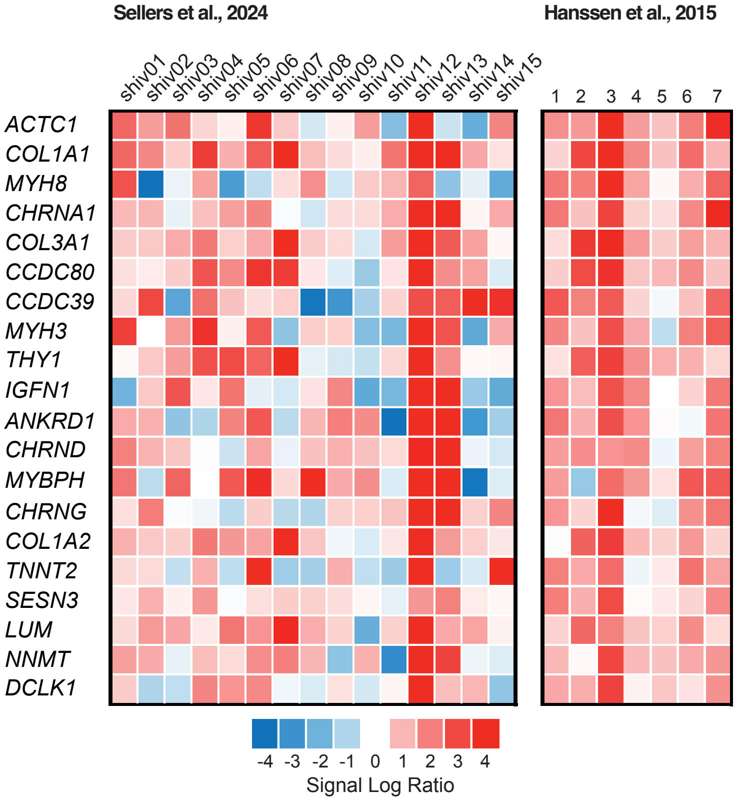


Heatmap of top 20 upregulated genes pre versus post cold acclimation in Hanssen et al., 2015, for all individuals of both Sellers et al., 2024 and Hanssen et al., 2015.
